# Characterizing the sub-second timescale strategies of fighting in zebrafish

**DOI:** 10.1101/208918

**Authors:** Andres Laan, Marta Iglesias-Julios, Gonzalo G. de Polavieja

## Abstract

Most animals fight by repeating complex stereotypic behaviors, yet the internal structure of these behaviors has rarely been dissected in detail. We characterized the internal structure of fighting behaviors by developing a machine learning pipeline that measures and classifies the behavior of individual unmarked animals on a sub-second timescale. This allowed us to quantify several previously hidden features of zebrafish fighting strategies. We found strong correlations between the velocity of the attacker and the defender indicating a dynamic matching of approach and avoidance efforts consistent with the cumulative assessment model of aggression. While velocity matching was ubiquitous, the spatial dynamics of attacks showed phase-specific differences. Contest phase attacks were characterized by a paradoxical side-ways attraction of the retreating animal towards the attacker suggesting that the defender combines avoidance maneuvers with display maneuvers. Post-resolution attacks lacked display-like features and the the defender was avoidance-focused. From the perspective of the winner, game theory modeling further suggested that highly energetically costly post resolution attacks occurred because the winner was trying to increase its relative dominance over the loser. Overall, the rich structure of zebrafish motor coordination during fighting indicates a greater complexity and layering of strategies than has previously been recognized.

## 1 Introduction

Animals fight by roaring [1], lunging [2], circling [3], head-waving [4], headbutting [5], biting [6], wrestling [4] and in a myriad of other ways [7]. While we have good theories and measurements about why an animal might start a fight with a fin-display and end the fight with mouth wrestling [8], we have have much less information about what exactly happens during a lunge, a circling display or a directed attack maneuver. Much of this may come down to the problem of measurement. It is comparatively easy to count the number of displays or time the duration of a wrestling bout, it is much harder to accurately measure a multi-dimensional signal like a threat display. We are therefore mostly confined to verbal descriptions of contest behaviors.

Yet the accurate measurement of within-behavior limb and body dynamics has been a source of rich insight in many other systems. When researchers were able to use high speed cameras to capture fly leg movements before they jumped to dodge a moving stimulus, they uncovered a sophisticated context-sensitive control system in what was previously believed to be a simple ballistic reflex [9]. Likewise, statistical descriptions of escape trajectories have given us convincing experimental evidence of protean behavior- a strategy where prey occasionally randomize their movement direction in order to reduce the degree to which their behavior can be predicted [10, 11]. The analysis of peregrine falcon attack trajectories has revealed a precise mathematical analogy between falcon prey capture and ballistic missile targeting which can only by uncovered through precision measurement [12].

The last two examples are particularly relevant for the study of aggression, because they illustrate cases where a complete understanding of strategic behaviors cannot be obtained unless we measure the dynamics occurring within elementary behaviors. Next we will highlight some outstanding issues in the study aggression which might similarly benefit from modern data capture methods.

As a first example, let us consider how much is known about attack maneuvers in zebrafish. During contests, zebrafish frequently engage in repeated attacks, where one animal performs a rapid directed movement towards another and the other animal sometimes responds with an avoidance/retreat maneuver [6]. What has remained unclear is the quantitative relationship between the attack maneuvers and the avoidance maneuvers. Does every attack induce an avoidance maneuver? Are the locomotor costs of an attack greater or smaller than the locomotor costs of a retreat?

One reason why answers to these questions matter concerns the theoretical interpretation of zebrafish fighting. Animal conflict is partly structured as a series of assessments of relative strength and different game theory models of assessment postulate a different relationship between individual activity levels and fitness costs. For example, WOA models postulate that individual acts of behavior induce fitness/energy costs only in the producer of the behavior [13] whereas SA [14] and CA [15] models allow the behavior of the producer to influence the fitness costs of the target of the behavior as well. Target fitness costs might occur because the target suffers contact injuries or alternatively, because it needs to perform costly avoidance maneuvers to avoid suffering the injuries. The relevance of the aforementioned factors to the interpretation of zebrafish contests is obvious: if attacks rarely end in contact and the cost of an avoidance maneuver is small compared to the cost of an attack, then a WOA model might potentially be a good description of zebrafish fighting. However, if each attack nearly always induces a costly avoidance maneuver in the target of the attack and in the absence of avoidance bodily contact typically occurs, then only the CA and SA model remain viable as descriptions of fighting.

A second utility to measuring the fine structure of fights stems from the potential to expand domains in which evolutionary game theory can be tested. As was already mentioned, it has been speculated that predator and prey interactions during escape maneuvers are best characterized as a game where the prey makes unpredictable maneuvers in order to avoid easy capture [10, 11, 16]. Similar games might unfold within the multitude of elementary interactions occurring during a fight. Analyzing games which occur on a fast time-scale might be of interest not only because it provides opportunities to expand the domain where game theory is applied, but also because payoffs obtained in low-level games will determine the payoffs of various strategies for the long time-scale assessment games in which they are embedded. A similar methodology has already shown promise in the analysis of schooling behavior [17].

In the last section of our results, we take a first step towards game-theoretical analysis of movement rules with a particular focus on the structure of resolution phase attacks. The literature on game theory and dyadic aggression is rich but appears to be primarily focused on the symmetrical/assessment phase of the conflict (see for example Chapter 2 of [18]). In addition to an assessment phase, zebrafish fights also have a structured postresolution phase, where both the winner and the looser engage in stereotypical behaviors. It has previously been shown that post-assessment behaviors either serve the function of chasing the looser out of the territory or maintaining dominance rank [19, 20, 21]. We show how insight from these studies can be used to formulate a model of zebrafish resolution phase attacks which provides a concise explanation of the main qualitative trends present in our data.

Thirdly, analysis of elementary aggressive interactions can shed light on multiple functions of a single behavior. When a boxer holds up his hands, it is with the dual purpose of being ready to both attack and defend. Likewise, a zebrafish attack may be shaped by multiple competing needs. The attack intensity may need to be moderated in order to avoid overcommitment to a single direction of assault which could be exploited by a responsive opponents. An attack might simultaneously carry out the function of damaging an opponent and signaling to it, or the dual functions of the attack may be somewhat separated in time. Without large-scale datasets, it is difficult to experimentally address these subtleties of multi-functionality and variability.

In order to measure zebrafish aggressive interactions at a level compatible with making progress on solving the above mentioned issues, it was imperative to create a new measurement system. We took inspiration from several pre-existing machine learning tools to create a system which allows for tracking and identifying unmarked animals as well as automatically annotating their behaviors [22, 23, 24]. The resulting system provides the user with trajectory data containing information about velocities, accelerations and relative positions of the fighting individuals as well as an automated ethogram which identifies the behavior performed by any animals at any given moment. Our set of tools provided us with the data necessary to examine aggressive behavioral interactions at the lowest possible level.

## 2 Methods

### 2.1 Staging of contests

The study used 68 male zebrafish of the AB strain approximately 1 year of age. All rooms are ventilated through a centralized HVAC system and are kept at controlled room temperature (25C), 50%–60% humidity. Fish holding rooms are kept under a 14-h light10-h dark cycle with a light intensity of 200 300 lux at the water surface. The density of the fish in the tanks was 10 fish/L and in a typical cage we had 20-25 animals. Our general feeding protocol consists of two types of live feeds, rotifers and Artemia nauplii, and a processed dry feed (Gemma Micro, Skretting, Spain). Depending on the fish age, the feeding frequency varies. In the months prior to experiments, the fish were fed Gemma500 feed and live de-capsulated Artemia once a day.

Our study involved staging contests between zebrafish. We adapted a procedure from [6] where a pair of males were removed from their home tanks and kept in visual but not olfactory isolation for a period between 24 to 48 hours. In a slight departure from the previous procedure, the fight was staged in an arena which was different from and larger than the arena used for prefight isolation, because we wanted to avoid the confounding influence of walls on swimming behavior which occurs too frequently in smaller arenas. The fight was staged in a uniform rectangular arena with dimensions 32-by-24-by-12 cm, slightly rounded corners and water depth of approximately 7 cm. Care was taken to ensure a lack of sharp illumination gradients in the tank so as to facilitate later tracking. Recordings began when the two animals were simultaneously poured from the isolation tank to the fight arena. A typical recording lasted for 1 hour and was continued for another hour in the rare cases where the fight appeared unresolved after 1 hours time. After the fight was terminated, both animals were returned to their home tanks. Video data was acquired at 20 frames per second using MATLAB standard functions.

### 2.2 Tracking

Code and trajectory data are available on request: https://goo.gl/eGCp3q Aggressive contests in zebrafish pose three challenges. First, fighting is a 3D process and the maneuvers are facilitated by deep waters, which induces appearance changes as the depth of the fish varies. Second, fish change their appearance not only due to varying depth but also due to color changes during the fight. Third, collisions are more frequent during fighting than during schooling. This motivated the use of a hybrid system where a new version of idTracker [22] (idtracker.ai, Romero-Ferrero, Bergomi et. al, in preparation) utilizing deep convolutional networks was used for tracking when the animals were not colliding, because of the greater expressive capacity of learned templates compared to the hand-engineered template of classical idTracker. When the pair of fish collided, a Gaussian mixture model was used to separate the colliding animals (as in [24]) and identity information was propagated into the collisions by using a greedy acceleration minimization principle along the trajectory with the constraint that identities of both trajectories at the start and end of the collision had to be matched with the predictions from idTracker (see [22] for an analogous algorithm for collision resolution).

The greedy acceleration minimization was implemented step by step. At each time step, two candidate coordinates (each representing the center of mass of a fish with unknown identity) originating from the GMM algorithm needed to be identified. We considered the identities of the coordinates at the previous two time steps to be fixed and then we calculated the absolute net linear acceleration along both trajectories for the two possible identity assignments. Whichever assignment resulted in the lower total acceleration was used for final identification and the cycle was repeated again and again until end of collision. During collisions, we had an identification accuracy of 98%.

### 2.3 Automated behavior classification and analysis

To improve data efficiency, we used a preprocessing method which was designed to reduce translational and rotational variance. For our four vectors at each time point, we transformed them into a new coordinate system where the zero was located at the joint center of mass of the pair of fish at time *t* — *K*. The x axis was aligned with a vector which pointed from fish 1 towards fish 2 at time *t* — *K*. All coordinates of the four vectors were converted into this coordinate system. After the preprocessing, the four processed vectors were then concatenated into a single vector and passed as input to the first layer of a standard multilayer perceptron with a ReLu hidden layer activation function, a cross entropy loss function. Using this preprocessing and a fairly small amount of annotated data (small as compared to the total corpus of data analyzed), we were able to train a perceptron with two hidden layers of size 250 neurons to have a test set accuracy of 95%.

All further analysis was done using custom-written MATLAB code. The forcemap technique was adapted from [25]. In order to avoid potential influences from walls, the symmetric phase forcemaps were analyzed only when both fish were further than 5 *cm* away from the nearest wall. During the asymmetric phase, the fish spent most of their time swimming very close to the wall, but we excluded the influence of corners on turning by removing all data when fish were closer than 5 *cm* to the nearest corner.

## 3 Results

We began our study by staging 34 contests between adult male zebrafish (see Methods for details of contest staging). In order to analyze the fine details of aggressive behavior in fish, we augmented the software idTracker [22] (which allows tracking and identification of unmarked animals) with a custom-written collision resolution system that enabled us to resolve the identities of the animals while they were colliding as well (see methods for details on this and all other machine learning procedures). The output of the tracking system was a time series of trajectories for both contestants (sampling rate 50 ms, 20 Hz). We annotated a small fraction of our video data to indicate when attacks were taking place. These annotations were in turn used to train a neural network which could detect the presence of attacks from trajectory data with 95% reliability. Our behavior classification system differed from similar systems [23] through the use of end-to-end deep learning on trajectory data, which allowed us to eliminate an intermediate feature engineering step. By combining several augmented and improved machine learning tools into a common pipeline, we thus created a machine learning software that automatically provided information about the movement, behavior and identity of each animal on a subsecond timescale (Figure 1).

**Figure 1:**
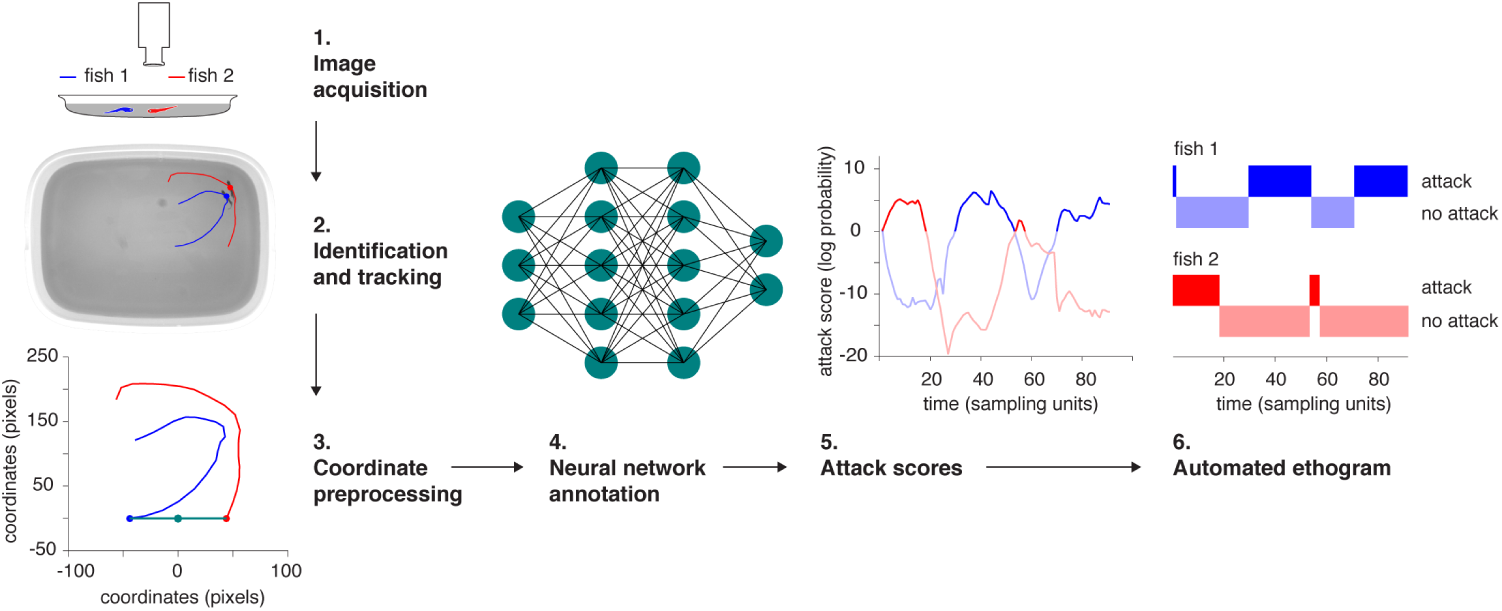
The computer vision pipeline. 1: the raw video. 2: unmarked animals after identification with idTracker and a short span of the trajectory of each animal overlaid. 3: preprocessing of a local chunk of trajectory for neural network analysis. 4: schematic of the neural network classifier which was trained to mimic human annotations. 5: a time series of attack scores for two animals as produced by the neural network classifier. High attack score values indicate a high internal confidence of the network that an attack is taking place. 6: an automatic ethogram calculated by thresholding the attack score.

### 3.1 Analysis of activity correlations and assessment models

As mentioned in the introduction, one of the focal points of our study was to achieve more precise measurements of locomotor activity in order to assess the relative costs of attacks and retreats. A necessary preliminary to studying activity correlations was to first characterize the large-scale patterns of aggressive behavior in our dataset. We found clear signs of aggression in 27 of the 34 staged contests. When we examined the attack rates of our contestants during individual contests, there was evidence that fights consisted of two distinct types of phases (Figure 2A, see also supplementary video S1). During what we called the symmetric phase (Figure 2A, 24-28 minutes), both individuals engaged in mutual attack behavior. During the asymmetric phase, mainly one individual performed attack behaviors (Figure 2a, 34-60 minutes). When a symmetric phase was present (N=15 fights), the most common pattern (N=8 of 15 fights) was for there to be a pre-fight phase with very few attacks (Figure 2A 0-22 minutes), followed by a symmetric phase, which was in turn followed by an asymmetric phase where only one individual engaged in attacks. It is thus likely that the symmetric phase is similar to the contest phase described in many other model systems of aggression, whereas our asymmetric phase resembles the resolution phase [21, 18].

**Figure 2:**
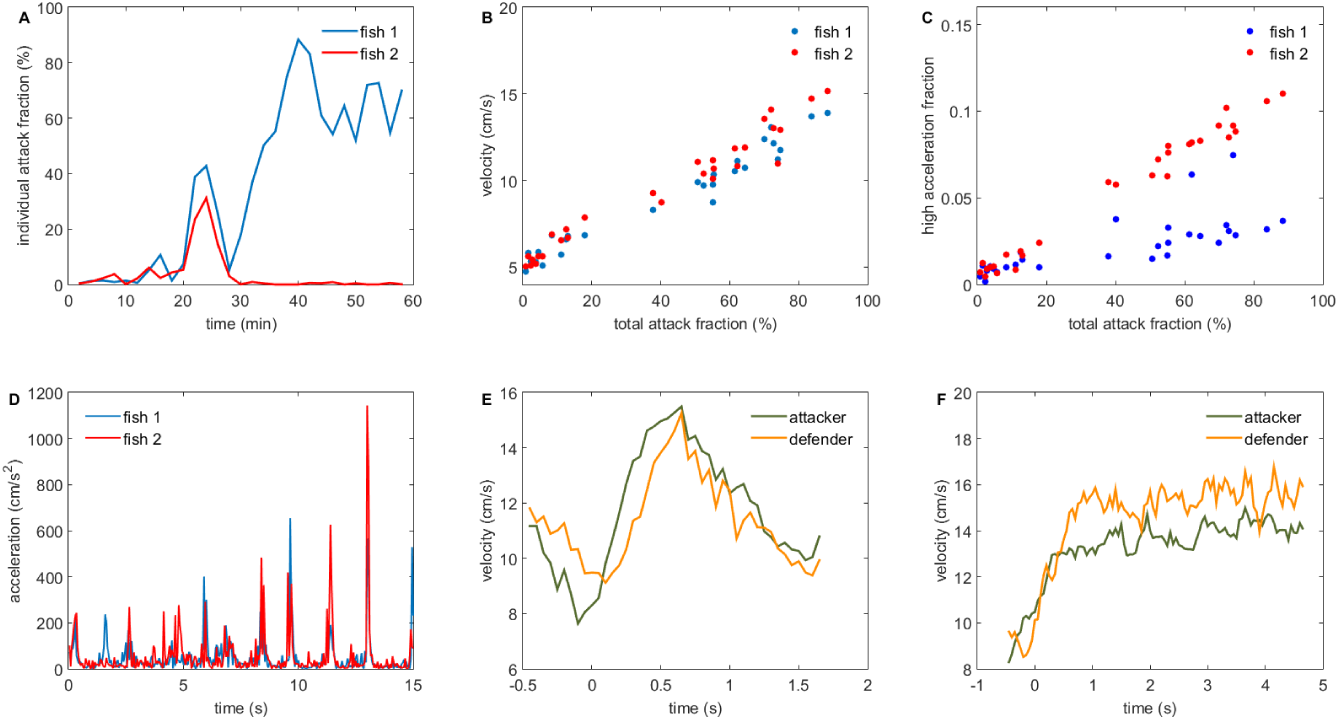
A kinematic characterization of a fight. A: The fraction of an animal’s time budget used in attacks (the individual attack fraction) over the course of the fight (blue and red curves mark the two different individuals here and elsewhere). Analysis is conducted in non-overlapping 2 minute time windows. Minutes 24-28 correspond to the symmetric phase. The asymmetric phase approximately spans the time from 34-60 minutes. B: correlation between total attack fraction (sum of individual attack fractions) and velocity. C: very high intensity acceleration bouts (*a* > 128 *cm/s*^2^) occur mostly during attacks. D: a time series of acceleration during the symmetric phase of the fight depict the occurrence of sudden acceleration bouts. E: the velocity waveforms of the attacker (green) and defender (orange) during an average attack (N=114) in the symmetric phase. Attacks begin at time 0. F: same as before but for the asymmetric phase (N=116).

However, not all fights followed the aforementioned progression. In some cases, the symmetric phase was not followed by an asymmetric phase and in other cases, an asymmetric phase both proceeded and followed the symmetric phase. Interestingly, the individual who was dominant before the symmetric phase was not necessarily the same one who engaged in attacks after the symmetric phase (see Figure S1 for example plots of fight progression in the more rare cases). In 12 fights, no symmetric phase was present and the only phase present was the asymmetric one. In the next sections, our analysis will focus on 14 of the 15 fights were the symmetric phase was present unless we state otherwise (with 1 fight excluded from analysis since its long duration posed a threat to animal welfare and had to be prematurely stopped).

We used the outputs of our machine learning pipeline to analyze coarse kinematic parameters of attacks, namely velocity and acceleration. We focused on these variables first since they may be regarded as an approximate individual level measure of energy expenditure [26]. Attacks in both the symmetric and asymmetric phase were associated with high velocities compared to the pre-fight phase. Pre-fight, the fish had an average speed of 5.3 ± 0.84 *cm/s* (N=13, mean ± standard deviation), which during the symmetric phase rose to 10.3 ± 1.9 *cm/s* (N=14) for the attacker and 10.9 ± 1.4 *cm/s* (N=14) for the defender. We note that here and elsewhere, the roles of attacker and defender were not fixed during the analysis of a fight but were calculated dynamically for each individual at each moment in time based on the outputs of our classifier. Fish swimming speed rose further during the asymmetric phase where the attacker attacked with speed 13.5 ± 1.6 *cm/s* (N=11) and the defender swam with velocity 14.0 ± 1.6 *cm/s* (N=11). As a further check of our analysis, we binned data from each fight into consecutive non-overlapping 2 minute long segments and calculated the average speed and the total percentage of time that attacks were occurring (the total attack fraction) during each time bin. There is a strong linear correlation between average movement speed and attack percentage (Figure 2B, *r* = 0.90 ± 0.06, N=28 individuals).

Fighting is associated not only with an increase in velocity but also with the occurrence of bursts of high acceleration (Figure 2D). As with speed, there was a strong correlation between the total attack fraction and the total fraction of time each animal spent performing acceleration bursts (Figure 2C, *r* = 0.88 ± 0.15, N=28).

From our analysis, it becomes clear that an attack induces a strong energetic cost not only for the attacker but also for the defender. This point is further reinforced if we time-align individual attacks and calculate the average velocity waveform for both the attacker and the defender during both the symmetric (Figure 2E) and the asymmetric (Figure 2F) phase. From Figure 2E,F it is apparent how attacks begin with an increase in the velocity of both the attacker and the defender. In fact, the locomotor costs for the defender are on average even higher than those for the attacker as defenders swim with a higher average speed in 20 out of the 25 conflict phases analyzed (p=0.004, two-tailed binomial test). These findings are compatible with the assumptions of the CA and SA models and violate the assumptions of WOA models.

As a control, we compared the inferences derived from our method with more established methods of analysis, which recommend disambiguating assessment strategies by studying the covariation between resource holding potential (fighting ability) and the duration of the contest phase (the symmetric phase in our terminology). The first step involved finding an indicator of resource holding potential. In our dataset, size was a statistically significant indicator of resource holding potential (RHP) as the larger animal ended up as the dominant individual in 20 out of 25 fights where we could identify a clear winner (p=0.002, one-tailed binomial test, analysis includes fights both with and without a symmetric phase). In fights where a symmetric phase was present, there was a statistically significant trend for large size differences to be associated with longer fights (r=-0.47, p=0.045, one-tailed t-test for Pearson correlation coefficient). A linear regression analysis of the effects of the sizes of both contestants yielded a model where the larger individual’s size had a negative effect on fight duration and the smaller individual size had a weaker but positive effect of fight duration, although the latter value was not statistically significantly different from zero (*C_large_* = —0.14,*C_small_* = 0.04, *P_large_* = 0.02, *P_small_* = 0.52).

A negative relationship between body mass difference and fight time is expected in all three models (WOA, CAM, SAM) [27]. A negative effect of larger individual body size on fight duration is incompatible with a WOA model of contest behavior. Our result, where the size of the larger individual has a stronger effect on fight times than the body size of the smaller individual is inconsistent with a pure sequential assessment game model, but as we show in a mathematical addendum, it is in principle consistent with the CAM model (see supplement, Mathematical analysis of the cumulative assessment model).

Overall, our analysis of resource holding potential and body size correlations yielded results which are consistent with the analysis of velocity correlations and thus indicates that our method might be of use as a substitute where the standard analysis is inapplicable or yields ambiguous results. A particularly interesting use case for our method might occur when fight times are found to be dependent only on the resource holding potential of the loser. Such a result has sometimes been interpreted as providing unequivocal support for WOA model [27], but other authors have held that this result would be compatible with CAM as well [4]. Our supplementary modeling agrees with the conclusions of [4] and thus motivates the need for further testing when loser-only fight time scaling relationships occur. Since our measurement methods allow discriminating between WOA and CAM, they may prove to be valuable for those further tests.

In addition to size, another weak predictor of fight outcome was color. We found that zebrafish exhibited a transient darkening which occurred specifically during the symmetrical contest phase (see Figure S2 and supplementary methods). On average, the symmetric fight phase was accompanied by an 8% ± 4% (N=28) darkening of appearance in both fighters and this transient largely disappeared irrespective of weather the fight ended with asymmetric chasing or not. The eventual looser tended to darken more than the winner. In 9 out of 10 fights, the eventual looser had a higher intensity change relative to pre-fight intensity than the eventual winner (p=0.02, 2-tailed binomial test). However, color change was a weak predictor of how the fight ended, since unequal changes in color were also associated with fights that ended without a clear way to determine the winner because chasing behavior was absent.

### 3.2 Analysis of movement rules

Zebrafish fight maneuvers have a complicated spatio-temporal structure, which may potentially contain useful information about strategic incentives. In order to analyze this structure, we utilized the tool of forcemaps which was originally developed to study cooperative movement coordination during schooling [28, 25, 17]. Forcemaps are useful for the study of aggression because they allow an easy visualization of the way fish influence each others movement during social interactions.

The central idea is to characterize the kinematic movements of a focal individual from its point of view. As a brief description of the procedure (see Methods for full details), during each moment in time, we transformed our trajectory data into a coordinate system where a focal fish is located at the center of the coordinate system and the y axis is aligned to the instantaneous velocity of the focal fish (see panel 3A,E and panel 4A,E; direct attention towards the orange and green dots). We then measured the distribution of locations for the partner of the focal fish in this new focal fish coordinate frame (see for example Figure 3B). We also calculated how the location of the partner fish influenced the tendency of the focal fish to turn and to speed up. As was the case with the analysis of velocity correlations, we separated our data into four categories depending on whether the focal fish was attacking or defending and whether the current phase of the fight was symmetric or asymmetric (in Figures 3 and 4 which follow, the colored fish on the inset images A,E always illustrates which fish is the focal fish for a given row of the figure).

**Figure 3:**
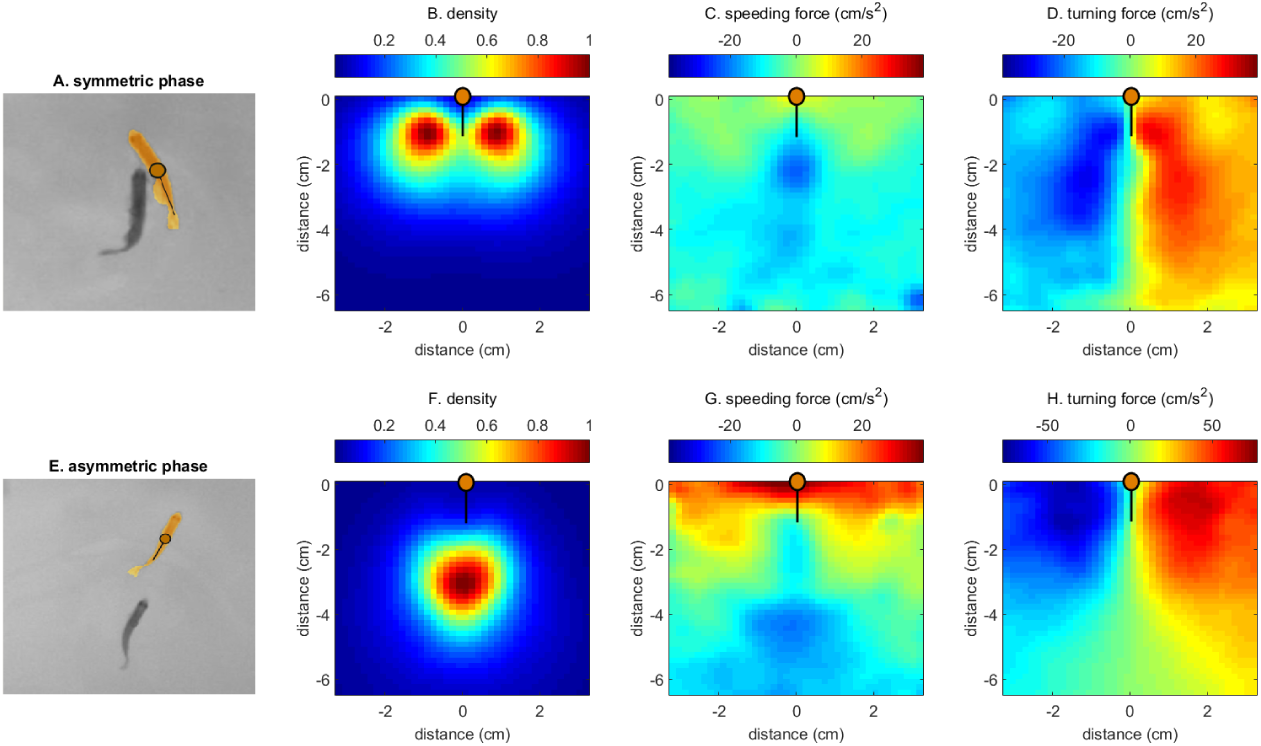
Forcemaps of the defender in the symmetric (top row) and asymmetric (bottom row) phase. Average forcemaps (N=14 fights, 230 000 time-points). The top inset image shows a typical configuration of the two fish during the symmetric phase (same for asymmetric phase at the bottom). In this figure, the focal fish is always located at the top middle of the map (orange dot, point 0,0) and its velocity vector is oriented up. A: a typical configuration of the two fish during the symmetric phase. B: a probability density map of the position of the attacker relative to the defender during the symmetric phase. Note that the negative numbers in the distance axis indicate that the attacker is behind the defender. C: the speeding force as a function of the relative location the attacker during the symmetric phase. Red color signals speeding up, blue color signals slowing down. D: the turning behavior of the defender as a function of the relative location of the attacker during the symmetric phase. Red colors signify turning to the right, blue colors signal turning to the left. E: a typical configuration of the two fish during the asymmetric phase. F-H: same as B-D for the asymmetric phase.

Previously, [6] had described zebrafish attacks as locomotion maneuvers where the attacker orients its body towards the defender and then swims rapidly towards it. The defender typically responds by swimming away from the attacker in a maneuver named retreat. Additionally, the attacker sometimes veers to one side or the other of the defender in order to deliver bites to the sides of the defender. Based on this description, we had four baseline expectations. First, the attacker is expected to be located behind the defender most of the time. Second, the attacker is expected to exhibit an acceleration response towards the defender if the defender is in front of the attacker. Third, the defender is expected to exhibit a repulsive speeding response when the attacker is behind it (the running away response). Fourth, when the attacker is located to one side (e.g. the right) of the defender, the defender was expected to turn towards the other side e.g the left) in order to dodge potential bites. To our surprise, all the hypothesis except the first turned out to be partly incorrect to varying degrees.

We begin by considering the relative positions of the defender and the attacker. The maps in Figure 3B,F depict the distribution of the locations of the attacker relative to the defender during attacks in the symmetric (3B) and asymmetric phase (3F). From the maps it is clear that as was anticipated, the attacker is typically located behind the defender. However, an interesting elaboration on the baseline hypothesis was the finding of different appearances of the maps between the two phases. During the symmetric phase, the attacker is located about half a body length behind the defender and is positioned to the left or to the right of the defender rather than staying directly behind it (Figure 3B). In contrast, during asymmetric phase attacks the attacker is typically located a whole body length behind the defender (Figure 3F). Since a sideways location of the attacker was expected to be associated with attempts of biting, the phase differences initially suggested a greater motivation on the side of the attacker towards delivering bites in the symmetric phase. While this view is likely to be partly correct, we will explore a further alternative explanation below.

Focusing on the speeding responses of the attacker next, we see that our initial expectation of seeing acceleration responses was again partly fulfilled. As can be seen in Figure 4C,G, if the defender is located far in front of the attacker, the attacker does indeed have a tendency to accelerate towards the defender as if it was attracted to the defender and was trying to chase it (the red and yellow areas at the top of the maps in Figure 4C,G). However, the expectation of pure attraction was violated at close range by the presence of repulsion zones (blue areas at the bottom of Figure 4C,G). If the attacker reached close to the defender, there was a tendency for the attacker to decelerate rather than accelerate. This deceleration response was present even if we removed periods of collision from the analysis (see figure S3C bottom panel). Therefore, the slowing down appeared to be a deliberate response by the attacker and not the consequence of some direct physical interaction like contact-driven repulsion.

**Figure 4:**
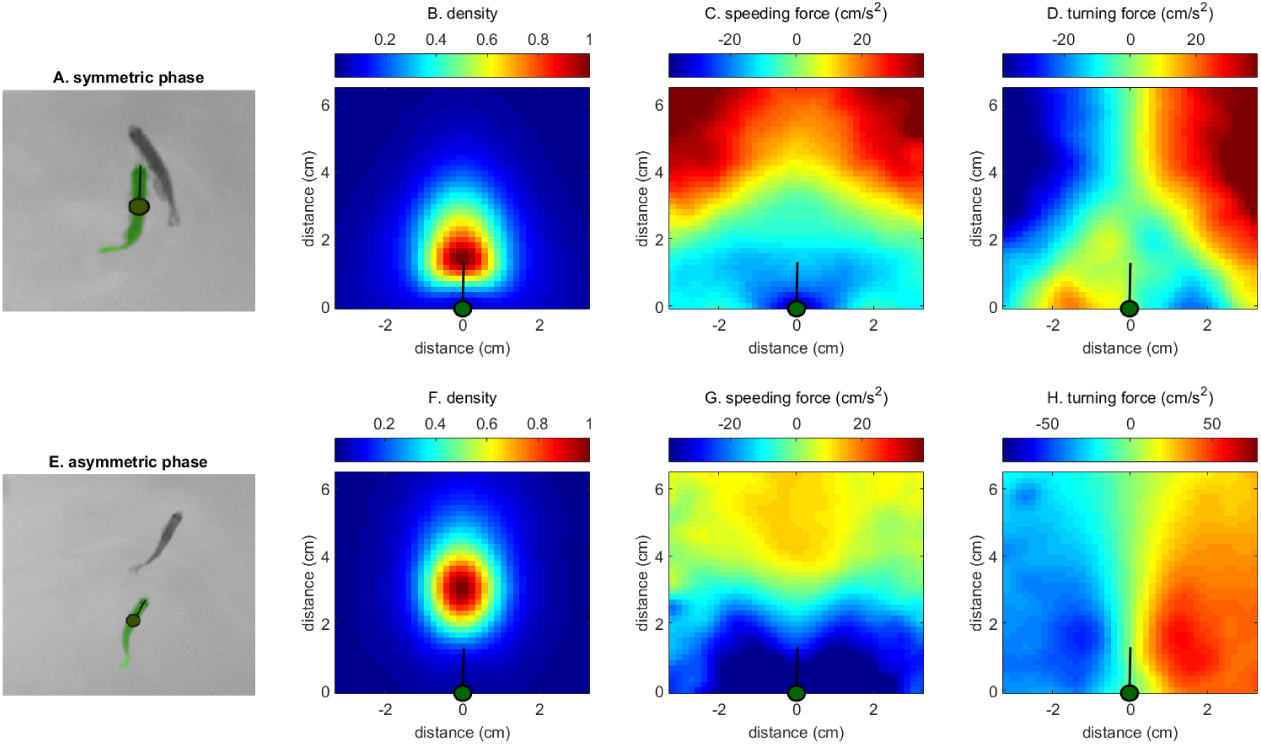
Forcemaps of the attacker in the symmetric (top) and asymmetric (bottom) phase. Average forcemaps (N=14 fights, 140 000 time-points). The focal fish for each row is marked in color on an inset image (A,E). On these forcemaps, the focal fish is always located at the bottom middle of the map (marked as a circle with a tail on the maps) and its velocity vector is oriented up. A: a typical configuration of the two fish during the symmetric phase. B: a probability density map of the position of the defender relative to the attacker during the symmetric phase. The positive numbers on the distance axis indicate that the defender is in front of the attacker. C: the speeding force as a function of the relative location the defender during the symmetric phase. Red color signals speeding up, blue color signals slowing down. D: the turning behavior of the attacker as a function of the relative location of the defender during the symmetric phase. Red colors signify turning to the right, blue colors signal turning to the left. E: a typical configuration of the two fish during the asymmetric phase. F-H: same as B-D for the asymmetric phase.

Overall the speeding map of the attacker was consistent with a strategy where the attacker tries to maintain a constant distance from the defender by speeding towards the defender if the defender was far and by slowing down and letting the defender escape if the defender was too close. This finding was at odds with our initial expectation that the primary goal of the attacks was to create bodily contact which would enable delivery of bites. However, we found evidence for such a distance maintaining strategy on the part of the defender was well. The speeding map of the defender in both phases showed acceleration responses (running away) when the attacker was too close (Figure 3C,G) and deceleration responses (permitting approach) when the attacker was far.

When we examined the turning maps of the attacker, it gave us further support for the idea that the attacker is using a distance maintaining strategy during the symmetric phase. As can be seen in Figure 4D, if the defender was far away, the attacker exhibited a turning response toward the defender. However, at close range, the turning response once again changed to a repulsive response (notice the flipped polarity at the bottom of the turning map when compared with the top in Figure 4D).

Examining the turning map of the attacker during the symmetric phase gave evidence of a more complex pattern than a simple distance-maintaining strategy on the part of the defender. We remind the reader of our initial expectation, which was for the defender to turn to the opposite side from the location of the attacker as an avoidance response. What we found was the exact opposite. If the attacker was to the left of the defender, the defender turned to the left towards the attacker.

The above response by the defender probably contributed to the stable maintenance of the T-like configuration (Figure 3A) which was often evident during the symmetric phase. Our initial hypothesis attributed the generation of the T configuration to efforts by the attacker to swim to this position so he could deliver biting attacks to the vulnerable sides of his opponent. Examination of social forces revealed the opposite was true. The defender appears to actively contribute to the maintenance of the T configuration by exhibiting a statistical tendency to turn towards the attacker thus exposing the sides of its body even further. Contact appears to be avoided because of a distance-maintaining strategy by the attacker instead. We emphasize again that this effect is not the result of physical contact forces because the maps have the same qualitative features even if we exclude from analysis the periods where the bodies of the two fish are in physical contact (see Figure S3).

We conclude that during symmetric phase attacks, at close range there is a paradoxical tendency for the defender to turn towards the attacker and for the attacker to avoid turning towards the body of the defender. Although we do occasionally observe contact and biting attacks between the contestants, the statistically typical behavior is towards mutual short-range avoidance.

The strongly peaked nature of the position histogram (Figure 3B) provides quantitative confirmation of the persistent nature of the T-like configuration. The persistence of the configuration in time in conjunction with the aforementioned evidence of repulsive interactions when the two fish are in the T-like configuration further supports our description of the T-configuration to be a metastable state. Such evidence also speaks against our initial hypothesis, which saw the T-like configuration as a non-persistent state arising either as a preamble to biting attacks or as a result of counter-maneuvers designed to avoid the bites. The balance of evidence seems to indicate that the T-configuration during symmetric phase attacks may instead be a ritualistic configuration which is maintained by mutual efforts and may thus function partly as a mutual display.

Crucially and consistently with the display hypothesis, the T configuration did not appear during post resolution attacks- presumably because display behaviors are superfluous after the winner has been resolved. As can be seen from Figure 3E,F, the typical configuration during post resolution attacks had the attacker located precisely behind the defender and the defender running away from the attacker in a straight line. This pattern is maintained due to the mutual presence of a distance maintaining strategy in the speeding maps (see above). Furthermore, if we look at the turning response of the defender in the region where the attacker is most likely to be located, we see a green zone of neutrality (Figure 3H, green triangle-like area at the bottom of the map) rather than an attractive turning response as was seen in the symmetric phase. The data therefore supports a notion where post-resolution decision maps help the defender simply to avoid the attacker rather than engaging in a more complex strategy.

One final feature of the forcemaps is worth noting. The maps can often be reliably calculated from data gathered only during individual fights. These individual maps have a structure that is qualitatively very similar to the plots we show in the main paper, where we pooled data across all 14 conflicts which exhibited a symmetric phase (see Figure S4,S5). These movement maps thus appear to be a very reliable feature of zebrafish aggression.

In addition to detailed analysis of attack bouts, we also found evidence for a heretofore undocumented type of aggressive maneuver which we termed the splash (Figure 5 top panels). We named the behavior splash after the characteristic wave-like ripple pattern which occurred on the water surface after the maneuver was performed. The splash behavior typically occurred as one zebrafish approached another. As the two zebrafish made contact (Figure 5B), one or both of them responded with a sudden acceleration maneuver (Figure 5A) which resulted in the orientations of the two fish being completely reversed and the two fish being propelled apart by a distance of a few body lengths or more (Figure 5C). Typically, a 180 degree change in orientation (Figure 5D) was completed in less than 50 ms. We believe the splash may be a crucial behavior which helps stabilize the previously mentioned display-like attacks and we will comment more on this hypothesis in the discussion.

**Figure 5:**
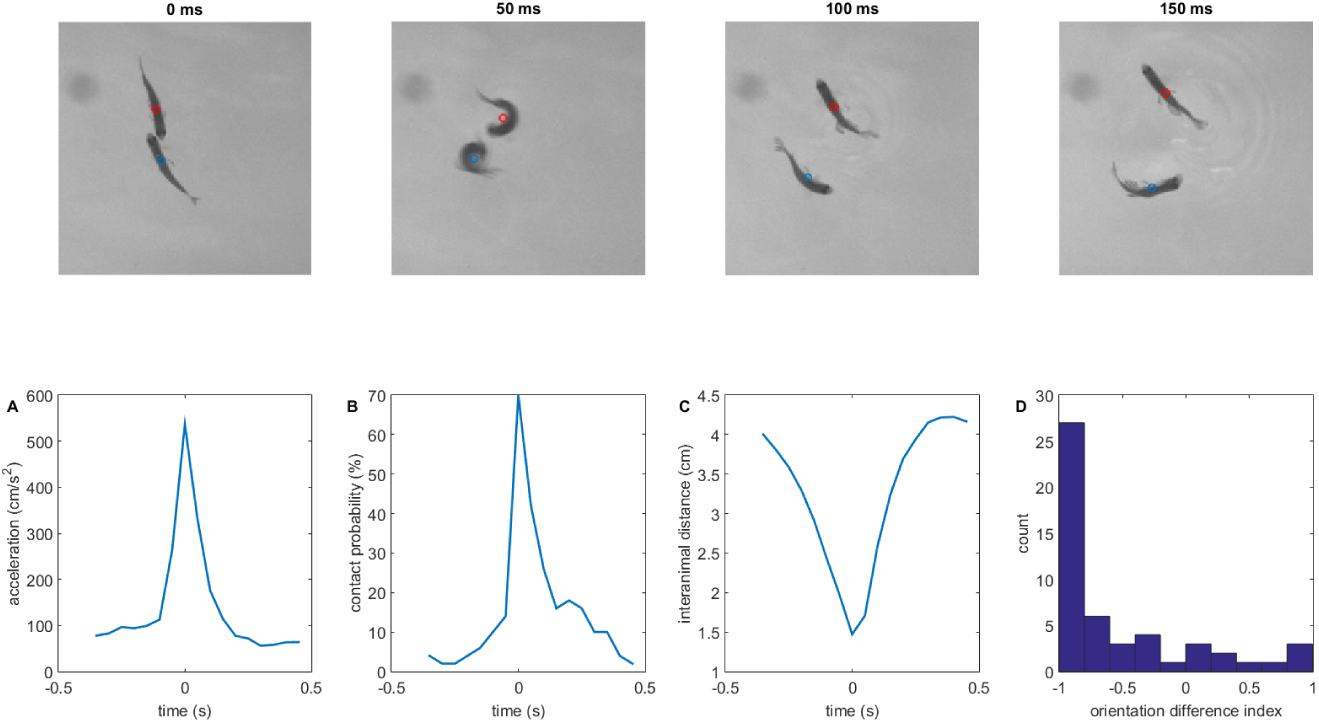
The splash behavior. Top panels: a time-lapse series of 4 consecutive frames (sampled 50 ms apart in time) during an example splash behavior. Bottom panels: A: the average acceleration of an individual during a splash. The splash takes place at time 0. B: The probability of contact during a splash. C: The average time evolution of inter-animal distance during a splash. D: A histogram of the orientation change index during 50 acts of splash. An index of −1 corresponds to a 180 degree change in orientation, an index of 1 corresponds to orientation being unchanged (see methods for details). For all plots, N=50 splash behaviors. Due to their short duration and comparatively rare occurrence, the splash behaviors were found by manual annotation.

### 3.3 Modeling of the asymmetric phase

We were motivated to seek a theoretical treatment of the post-conflict phase by the observation that both the winner and the looser engage in a contact-free but high-velocity chases, which are costly for both the winner and the looser. Why does the winner use a strategy which results in a waste energy on attacks even after it has established its dominance in the symmetric phase? Based on previous studies (see [19, 20, 21] as well as introduction) we hypothesized that the winner implements a strategy which simultaneously damages the loser without risking the possible loss of dominance.

We can therefore think of the post-resolution fight as a zero-sum game, where the reward of the winner is *r_w_* = *C_l_* — *C_w_* and the reward of the looser is *r_l_* = *C_w_* — *C_l_*(*C_w_* and *C_l_* designate the costs incurred by the looser and the winner respectively). What are the possible ways that two fish can impose costs on one another? Zebrafish incur costs either through rapid swimming or by receiving bites from the opponent. During the post-resolution phase, the dominant engages in rapid swimming with an approximately constant velocity (Figure 2F) while staying a constant distance away from the subordinate (Figure 3F). Our analysis must explain why the dominant never accelerates enough to touch and deliver bites to the opponent, which would certainly help to selectively reduce subordinate fitness.

In order to maintain a stable velocity *v*, the winner fish must generate a force *F*(*v*) which carries a cost *C*(*F*). If the looser also swims with velocity v, it will also incur a cost *C*(*F*). If the winner engages in biting, it will deliver to the looser an additional cost *C_b_*. However, it is reasonable to expect that the opportunity to deliver bites at velocity *v* will not come without a cost to the attacker as well. It must produce extra force in order to generate some pressure between his own mouth and the body of the looser. In addition, extra energy may be needed for moving the jaws and potentially suffering a less streamlined posture because of the bending needed to deliver the bites. The extra force needed *δF* will induce a greater cost of *C*(*F* + *δF*), while the looser incurs a cost of only *C*(*F*).

For real fish, the functions *F*(*v*) and *C*(*F*) are obviously not completely generic. *F*(*v*) is monotonically increasing, because higher velocities require higher forces in order to overcome increased drag. The function *C*(*F*) is likely to be not only monotonically increasing, but also convex, since maintaining higher forces requires recruitment of more energetically inefficient muscle groups [29]. For convex functions, Δ*C* = *C*(*F* + *δF*) – *C*(*F*) is an increasing function of *F* as well. At equilibrium, it must be the case that the winner cannot increase his reward by switching from steady chasing at velocity *v_e_* and force *F*(*v_e_*) to a biting attack at velocity *v_e_* and force *F* + *δF*. Therefore, at equilibrium, Δ*C*(*F*) = *C_b_*. Since Δ*C*(*F*) is an increasing function of *F*, it is a mathematical inevitability that there exists a value of *F* high enough for this to condition to be true. Another solution to the model involves limits on the range of possible values of F. If the value F has a biological maximum (*F_max_*) which is smaller than the value of F where biting costs become equal to attack costs, then the equilibrium value *v_eq_* is given by *F*(*v_eq_*) = *F_max_* — *δF*. In both cases, the equilibrium involves a stable value of *v_eq_*.

From our theory, we conclude that it is necessary for the looser fish to maintain the high velocity *v_eq_* as well because otherwise biting attacks become profitable for the winner and the looser will further descend in relative capacity. The winner in turn must maintain a high velocity and a close distance from its opponent or else the looser may respond by slowing down since the dominant is too far away to attack. The caudal deceleration zones apparent in the speeding map of the defender in the asymmetric phase (Figure 3G) may well be a mark of such strategic responses. The analysis thus indicates a plausible link between game theory equilibria, well-known features of fish biomechanics/physiology and the observed long duration chasing which often concludes zebrafish fights.

## 4 Discussion

We have introduced a machine vision pipeline for the study of aggression in zebrafish which allows both automated identification and tracking (see Methods) of unmarked animals by use of idTracker [22] as well as individual-level automated classification of ethologically relevant behaviors on a sub-second timescale (compare with the lack of identification and end-to-end deep learning in [23, 24]).The pipeline allows for reduced human workload by elimination of the marking stage and the annotation stage as well as reducing the need for controls comparing marked and unmarked animals. Our methods also have the additional advantage of allowing for some parallelization. Though we focused here on experiments in large arenas to avoid confounding influence of walls on our analysis, it is possible to fit up to 4 smaller fight arenas into the field of view of our camera. The tracking can also be done in parallel without modifications to the code. Hence, it is feasible for certain experiments to increase the setup throughput by a factor of 4 if needed.

In our work, we have also demonstrated how the ability to gather high resolution trajectory data can be of aid in the process of deciding which of the many assessment models provides the best description of the fight. For example, we observed a strong correlation between the velocity of the attacker and the defender during individual attacks. The observation of strong mutual correlation in activity levels during individual acts of behavior gives evidence that in zebrafish, approximately equal locomotor costs are borne by both the producer of the attack as well as its target. This observation rules out WOA models of contests as good descriptions of zebrafish aggression. We reach the conclusion because these models posit individual behavioral acts to have an effect on the energy budget of only the one who is producing the signal and not on the target of the signal [13]- a hypothesis clearly violated in our data.

Based on our results, the standard sequential assessment game also appears ill-suited as a description of zebrafish aggression, because we were unable to detect statistically significant positive relationship between the fight time and the RHP of the looser [27]. Having ruled out both the self-assessment and the sequential assessment models, we were left by elimination with the cumulative assessment game as the only suitable description of zebrafish fighting and our supplementary modeling supported this conclusion as well.

As stated before, we speculate that our technology may become a valuable complement to the current standard methodology of game theory model testing in two contexts. First, as others have argued [4] and as we showed in our supplementary modeling, the distinction between the CAM and WOA in terms of fight time scaling relationships is not as clear-cut as is sometimes stated [27, 30]. In such cases, the use of machine learning tools to infer activity budgets and correlations from video data may become a valuable complement to the standard toolkit as it will occasionally allow resolution of the ambiguities. Secondly, since our analysis does not require knowledge of the resource holding potential, it can be used in cases where the RHP is unknown or RHP differences are small.

Beyond the falsification of game theory models, analysis of trajectory level data also proved useful in clarifying the nature of certain behaviors. In the beginning we believed that the primary function of attacks in the symmetric phase was to maneuver the attacker into a position where he might be able to elicit further damage trough direct contact and biting. We were surprised to find in our forcemaps that rather than avoiding such attacks, the defender had a statistical tendency to turn its flank towards such attacks. Even more surprisingly, the attacker had a tendency to incompletely exploit the resulting vulnerable configuration as evidenced by the presence of weak repulsion zones in the turning rule of the attacker. The willingness of the defender to expose its flank may thus be at least partly a display behavior intended to signal its ability to maneuver and/or withstand damage.

The potential risk of the display may be mitigated by the opportunity to engage in the splash behavior. The splash behavior may enable one fish to halt or perturb the approach of another as it is becoming too dangerous. In support of this, notice how the splash is usually deployed right as the attacker is making first contact with the defender (Figure 5B). The potential option to engage in the splash behavior may also mitigate the risk associated with engaging in the display-like attacks which leads the defender into a vulnerable configuration. The vulnerable configuration which occurs during the displays may be stabilized because the attacker knows that any attempt to exploit the vulnerability can be countered with a splash maneuver by the defender.

One of the novelties in our paper was the introduction of movement rules [25] to the analysis of contests. The ability to quantify the fine structure of aggressive attacks trough movement rules is useful not only for the insight it provides about typical fighting tactics, but also because it enables quantification of change in those tactics. There is now much evidence for the role of cognition and learning in shaping animal fighting ability [31]. Fighting ability of animals changes with experience [32, 33], but exactly how experience makes fighters more competent and skillful has not always been clear from the studies. It may be that changes in fine motor dynamics play an important role and our measurement toolbox could be helpful in clarifying some of these unresolved issues. For example, evidence from sticklebacks has established a role for learning in the development of displays [34]. If the same is true for zebrafish, then there is an expectation that early in development, contest phase attacks might lack some of the display-like features we see an adults. The forcemap technique we have introduced could be straightforwardly applied to address this hypothesis, which might prove more difficult to test with traditional methods.

Finally, we hope that the study of trajectory level data will open up a new frontier in he study of strategic conflict. With the ability to record high resolution data, we may be able to get a better handle on the biomechanical determinants [17] of movement during contests. This may finally allow us to study the long-ago stated goal of examining not just how displays are used, but what factors determine the form and the fine dynamics of the displays as well [35]. Or in other words, we may eventually be able to study the movement subgames taking place within the larger assessment games. We took a small step in that direction by explaining the qualitative patterns of locomotion during the chasing phase through a game theory analysis, but there is also a clear need for better theoretical methods to analyze the extended games which occur when acceleration decisions influence inter-individual distances over time. The recent merging of techniques from game theory and deep reinforcement learning represents a promising avenue for further research in this regard. In particular, the use of self play, which has allowed humanoid robots to teach each other wrestling in an unsupervised way, is a technology which should be immediately applicable to the study of fish aggression as well [36].

## Author Contributions

Designed project: MJ AL GP. Performed the experiments: MJ AL. Wrote code: AL. Analyzed the data: MJ AL GP. Obtained models: MJ AL. Contributed to the writing of the manuscript: MJ AL GP.

## Conflict of interest statement

The authors declare no conflict of interest exists

## Acknowledgements

We wish to thank Francisco Romero-Ferrero and Mattia Bergomi for sharing an early version of idTracker.ai as well as for advice on how to best utilize the software. We thank Hanna Kokko and members of the Collective Behavior Lab for discussions.

## Funding

We acknowledge funding from the Champalimaud Foundation (to GdP) and from Fundaçao para a Ciência e Tecnologia PTDC/NEU-SCC/0948/2014 (to GdP) and FCT fellowships (to AL and MJ).

## Supplementary video link

https://drive.google.com/open?id=0Bz9JP7PG5u41S1phejlyWU5Wb3M

